# Transcriptomic signatures predict regulators of drug synergy and clinical regimen efficacy against Tuberculosis

**DOI:** 10.1101/800334

**Authors:** Shuyi Ma, Suraj Jaipalli, Jonah Larkins-Ford, Jenny Lohmiller, Bree B. Aldridge, David R. Sherman, Sriram Chandrasekaran

**Affiliations:** Center for Global Infectious Disease Research, Seattle Children’s Research Institute, Seattle WA, USA; Pathobiology Program, Department of Global Health, University of Washington, Seattle, WA, USA; Department of Biomedical Engineering, University of Michigan, Ann Arbor, MI, USA; Department of Molecular Biology and Microbiology, Tufts University School of Medicine, Boston, MA, USA; Laboratory of Systems Pharmacology, Harvard Medical School, Boston, MA, USA; Sackler School of Graduate Biomedical Sciences, Tufts University School of Medicine, Boston, MA, USA; Department of Biomedical Engineering, Tufts University School of Engineering, Medford, MA, USA; Center for Computational Medicine and Bioinformatics, University of Michigan, Ann Arbor, MI, USA; Microbiology Department, University of Washington, Seattle, WA, USA

**Keywords:** Tuberculosis, drug combinations, transcription factors, drug synergy, transcriptomics

## Abstract

The rapid spread of multi-drug resistant strains has created a pressing need for new drug regimens to treat tuberculosis (TB), which kills 1.8 million people each year. Identifying new regimens has been challenging due to the slow growth of the pathogen *M. tuberculosis* (MTB), coupled with large number of possible drug combinations. Here we present a computational model (INDIGO-MTB) that identified synergistic regimens featuring existing and emerging anti-TB drugs after screening *in silico* over 1 million potential drug combinations using MTB drug transcriptomic profiles. INDIGO-MTB further predicted the gene Rv1353c as a key transcriptional regulator of multiple drug interactions, and we confirmed experimentally that Rv1353c up-regulation reduces the antagonism of the bedaquiline-streptomycin combination. Retrospective analysis of 57 clinical trials of TB regimens using INDIGO-MTB revealed that synergistic combinations were significantly more efficacious than antagonistic combinations (p-value = 1 × 10^−4^) based on the percentage of patients with negative sputum cultures after 8 weeks of treatment. Our study establishes a framework for rapid assessment of TB drug combinations and is also applicable to other bacterial pathogens.

**IMPORTANCE:** Multi-drug combination therapy is an important strategy for treating tuberculosis, the world’s deadliest bacterial infection. Long treatment durations and growing rates of drug resistance have created an urgent need for new approaches to prioritize effective drug regimens. Hence, we developed a computational model called INDIGO-MTB, which identifies synergistic drug regimens from an immense set of possible drug combinations using pathogen response transcriptome elicited by individual drugs. Although the underlying input data for INDIGO-MTB was generated under *in vitro* broth culture conditions, the predictions from INDIGO-MTB correlated significantly with *in vivo* drug regimen efficacy from clinical trials. INDIGO-MTB also identified the transcription factor Rv1353c as a regulator of multiple drug interaction outcomes, which could be targeted for rationally enhancing drug synergy.

## INTRODUCTION

Tuberculosis (TB) is a global health threat of staggering proportions, taking a human life every 30 seconds (1). To ensure adequate treatment and combat onset of resistance, TB patients receive multidrug therapy. However, the frontline regimen of four drugs and six months treatment has not changed in 50 years, and resistance is spreading. In response, experts have called for entirely new regimens to combat the TB pandemic (2). While some new anti-TB agents are beginning to emerge (3), optimizing individual agents into effective regimens remains a significant challenge.

At present, combinations are designed and tested empirically, driven in part by clinical intuition. A standard approach to evaluate drug interactions experimentally utilizes checkerboard assays, which involves exposing the pathogen to different dose combinations of constituent drugs in a regimen. New approaches have been developed to increase throughput of checkerboard assays, either by reducing the number of doses required or by using computational optimization to find optimal doses (4-6).

Even with these developments, the enormous and expanding number of potential drug combinations renders regimen optimization by comprehensive experimental testing infeasible. The 28 drugs used to treat TB (7-10) could be assembled into nearly 24,000 different 3- or 4-drug combinations. Adding just two new agents to that list increases the number of different combinations to almost 32,000. Thus, there is a need for high-throughput approaches that can prioritize new drug combinations based on data generated from individual drugs. For example, a feedback-based approach was recently used to determine the optimal dosing of multi-drug regimens (4, 5). However, this approach still requires hundreds of dose-specific measurements for training the algorithm, all of which must be re-done whenever a new agent is under consideration. Computational tools such as metabolic modeling, kinetic modeling, and statistical modeling (11-13) have limited power in this context because direct targets are not known for many compounds. Existing approaches are also limited in the scale at which potential combinations could be evaluated computationally — currently around hundreds. Furthermore, empirical approaches based on drug similarity (or dissimilarity) are less effective in predicting interaction outcomes for new drugs classes, and they also lack a model for antagonism (14). Drugs with similar targets can have both synergistic and antagonistic outcomes (14).

To address this challenge, here we extend an *in silico* tool that we recently created —<u>In</u>ferring <u>D</u>rug Interactions using chemo-<u>G</u>enomics and <u>O</u>rthology, (INDIGO) (14)— to predict synergy/antagonism in combinations of two or more drugs. The original INDIGO model used chemogenomic profiling data under exposure to individual drugs (15, 16) as input data to identify drug-response genes (14). The scientific premise underlying INDIGO is that drug synergy and antagonism arise because of coordinated, systems-level molecular changes involving multiple cellular processes. Importantly, INDIGO can learn patterns from known drug interactions, which can then be used to forecast outcomes for new drugs and conditions. INDIGO can thus provide insights on underlying mechanism of drug interactions in an unbiased fashion. INDIGO can assess millions of combination regimens without requiring information about the drug target or mode of action. Once an optimal drug regimen can be determined using INDIGO, the dose regimes could be further optimized using feedback-based dose optimization techniques (4, 5).

The goal of this study is to identify antibiotic combinations that are most promising for TB drug development. We have adapted INDIGO to make use of transcriptomics data to identify drug-response genes, which are more widely available than chemogenomics data for most non-model organisms, including *Mycobacterium tuberculosis* (MTB) (**Figure 1**). We then harness a large compendium of publicly available and in-house generated transcriptomics data to show that INDIGO can successfully estimate drug interactions in MTB. We further integrate INDIGO with known MTB gene regulatory networks to identify transcription factors (TFs) that influence the extent of synergy between drugs. False positives and outliers from our model represent existing knowledge gaps and can inform future drug interaction experiments. The significant correlation of INDIGO-MTB predictions with both *in vitro* validations of novel predictions and *in vivo* efficacy metrics from clinical trials indicate that the INDIGO-MTB model has great promise for selecting novel TB drug regimens. INDIGO-MTB further provides unbiased insights on underlying cellular processes that influence drug interactions.

**Figure 1.**
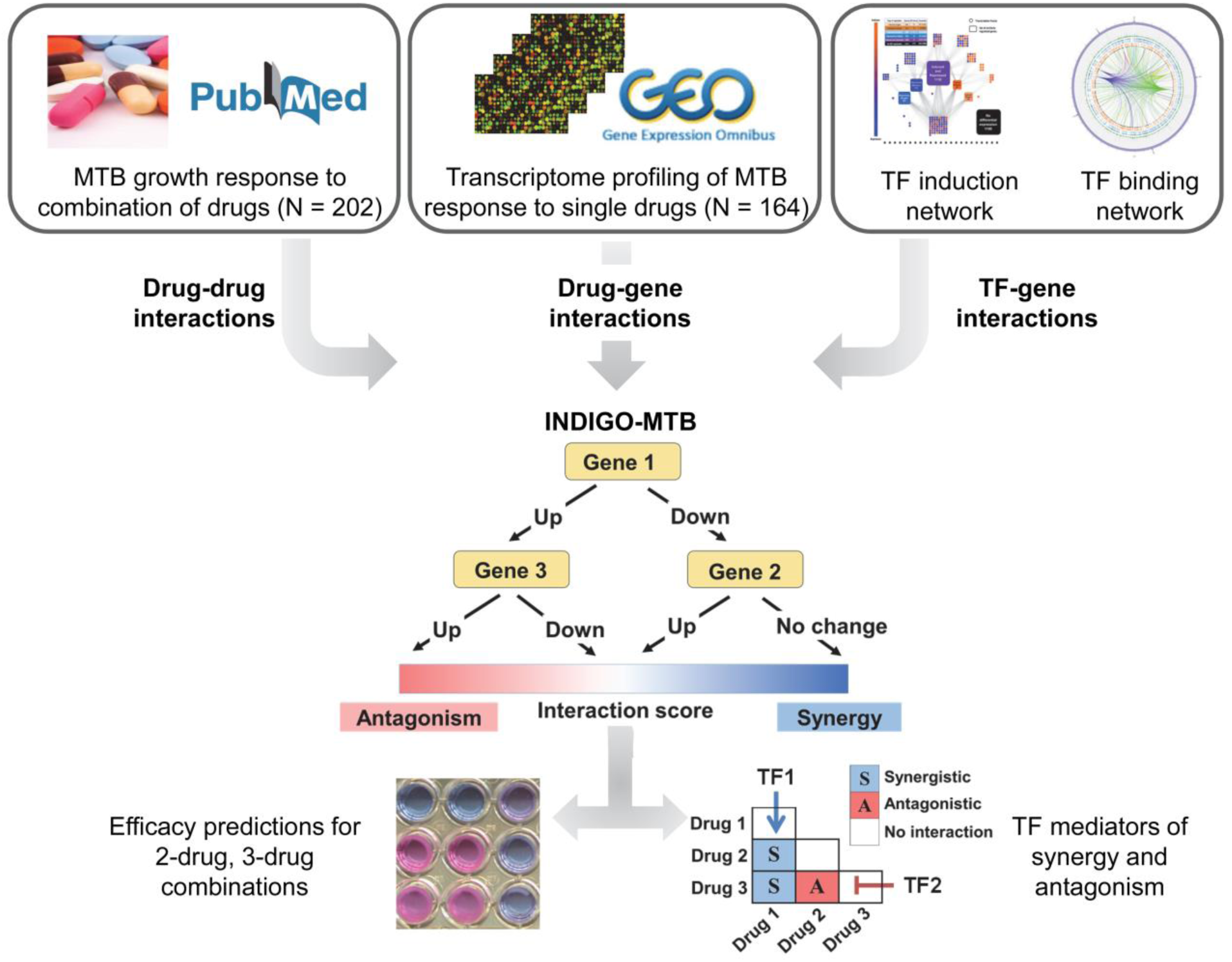
Schematic of INDIGO-MTB. INDIGO uses drug-gene associations inferred from transcriptomic data and experimentally measured drug-drug interactions as inputs to train a computational model that can infer interactions between new combinations of drugs. It does this by learning patterns in the drug-gene associations that are correlated with synergy and antagonism. In the example above, MTB upregulation of both gene 1 and gene 3 in response to the drugs measured in monotherapy is predictive of antagonism when the drugs are combined. By perturbing individual genes and known targets of Transcription Factors (TFs) in the model, we can infer the impact of individual gene and TF activity respectively on drug interactions and subsequently engineer interaction outcomes.

## RESULTS

### Construction of INDIGO-MTB model from drug response transcriptomes

The INDIGO approach requires a list of drug-gene associations and known drug-drug interaction data as input for building a chemical-genetic model of drug interactions. A gene is assumed to have a chemical-genetic association with a drug if a change in its expression leads to a statistically significant alteration in sensitivity to the drug of interest. A drug-gene association network is created by integrating chemogenomic profiling data from hundreds of drugs. This static network is then converted into a predictive model by leveraging the powerful statistical learning tool, Random Forest (17). This algorithm builds decision-trees using genes in the drug-gene association network and identifies those that are predictive of drug interaction outcomes using a training data set. The training data comprises known drug interactions. This trained network model can be used to forecast interactions for novel drug combinations (**Figure 1, Figure S1**).

While in the prior study, drug-gene associations were obtained from chemogenomic profiles of *E. coli*, these comprehensive gene deletion/drug response data are difficult to generate experimentally for most pathogens. We hence hypothesized that transcriptomics data, which quantifies the responses of every gene to a given perturbation, could provide a readily available alternate resource for analysis. This solution could circumvent the limitation that chemogenomic data are not available for most pathogens, including MTB. Generating gene expression data for response to monotherapy drug exposure is straightforward, and there are already publicly available transcriptomic profiles for many anti-TB agents.

We compiled transcriptome data profiling MTB response to different compounds and metabolic perturbations from the literature. We augmented this compendium by generating MTB transcriptomic response profiles for emerging TB agents (Methods, **Table S1A**). In addition to these transcriptomic data, we also used chemogenomics data from *Escherichia coli* (16), with *E. coli* genes matched to corresponding orthologous genes in the MTB genome. Our prior study showed that INDIGO can infer interactions in MTB with significant accuracy using orthologous gene mapping (correlation R = 0.54; p-value = 0.006). This was based on the observation that genes predictive of drug-drug interactions were surprisingly conserved between *E. coli* and MTB. In cases where multiple datasets profiled the same compound, we prioritized data from MTB profiled with the latest transcriptomics technology whenever possible. We normalized this drug response compendium using the ComBat algorithm (18) to account for inter-study and technology-specific (i.e. microarray, RNAseq) variation in transcriptomics data (Methods). Overall, this compendium contains data for 164 compounds and 65 metabolic perturbations (see **Table S1A** for full list)(12, 19, 20).

To train INDIGO-MTB, we compiled drug interaction values in MTB for 202 drug combination regimens from the literature, featuring compounds with available chemogenomic or transcriptomic profiles (**Table S1B**). The drugs in the training set consist of well-established anti-TB drugs, including rifampicin (RIF), isoniazid (INH), streptomycin (STM), several fluoroquinolones, as well as new drugs such as bedaquiline (BDQ). The extent of interaction between drugs was quantified in these studies by the standard Fractional Inhibitory Concentration (FIC) index (21), or the DiaMOND interaction score (6). In both of these metrics, synergy implies that the same amount of growth inhibition is achieved with a lower dose when both drugs are combined. We used statistical data normalization to combine these datasets, similar to our approach for combining transcriptomics data from various studies and platforms (Methods). This allowed us to account for the new technology-specific variation in drug interaction score distribution. If separate studies in literature provided conflicting interaction scores for a drug combination, we included both values to incorporate this experimental uncertainty into the model.

### Experimental validation of INDIGO-MTB model

The INDIGO-MTB model trained on these drug interaction data was used to infer interaction outcomes for new drugs and regimens. Given that our compendium has 164 compounds and 65 perturbations, INDIGO-MTB estimated all 26,106 potential pairwise interactions and all 1,975,354 potential three-way interactions. **Table S1C** shows the entire list of pairwise combinations and interaction scores.

We observed striking associations between specific compounds and interactions that were highly synergistic or antagonistic. In particular, combinations containing the drugs chlorpromazine and verapamil were highly enriched for synergistic interactions; 77% of chlorpromazine-containing combinations and 80% of verapamil-containing combinations were found to interact synergistically (FIC < 0.9) (**Figure S2**). Verapamil is an efflux pump inhibitor that influences membrane potential (22) and has been previously been shown to potentiate the activity of several anti-TB drugs (23-25). In contrast, all pairwise combinations featuring sutezolid were found to be antagonistic.

Previous work had found combinations of bacteriostatic drugs paired with bactericidal drugs were likely to be antagonistic against *E. coli* (26). INDIGO-MTB uncovered a similar trend in the MTB drug interactions; combinations featuring a bacteriostatic drug and a bactericidal drug had significantly more antagonistic interaction scores than combinations featuring only bacteriostatic drugs (p < 10^−12^). Interestingly, combinations featuring only bactericidal drugs also had significantly more antagonistic interaction scores than combinations featuring only bacteriostatic drugs (p < 10^−12^) (**Figure S2**).

To evaluate the accuracy of INDIGO-MTB, we experimentally measured interactions between a set of two-drug and three-drug combinations, and we compared these measurements against the interaction scores from INDIGO-MTB. The compounds featured in the tested combinations are all FDA-approved agents that have diverse mechanisms of action, and are either part of current first-and second-line TB therapy, or have been previously studied for their anti-tubercular activity. The interaction outcomes for the test set combinations spanned the entire range of INDIGO-MTB predicted interaction scores, enabling a rigorous assessment of INDIGO-MTB (**Figure 2A**, **Figure S3**). We quantified the interaction outcome either by traditional checkerboard assays or the high-throughput DiaMOND method for three-way combinations (Methods). Given the diverse methodologies used in literature for measuring drug interactions, we included combinations frequently measured in prior literature involving INH, RIF and STM as reference combinations in our test set. In addition, among the test set combinations, 10 combinations involved pairwise subsets of three-way combinations that were measured using DiaMOND methodology to infer 3-way interactions. Overall, among the 36 combinations in the experimental validation set, 24 combinations were completely “novel”, i.e. never seen by INDIGO. The sample size (N = 24 combinations) for the test set chosen for experimental validation is sufficiently powered to significantly assess the accuracy of INDIGO’s correlation with the experimental data (Methods).

**Figure 2.**
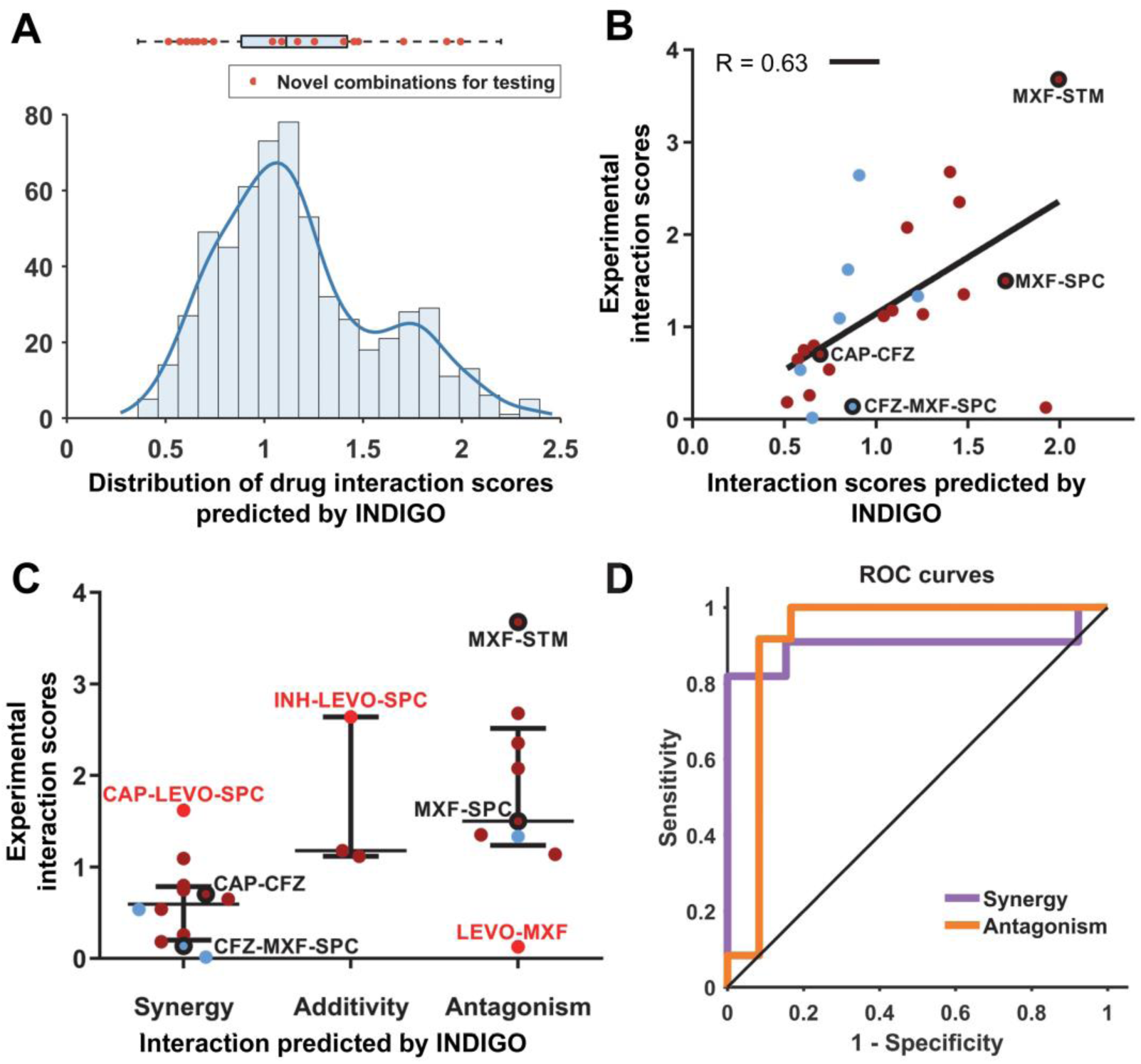
INDIGO-MTB accurately predicts novel drug interactions. **(A)** Drug combinations chosen for experimental testing span the entire range of drug interaction predictions by INDIGO. The histogram and box plot above it show the distribution of pairwise drug interaction scores for the 35 high interest TB agents (the edges of the box plot demarcate the 25^th^ and 75^th^ percentile, and the dashed lines extend between the 1^st^ and 99^th^ percentile). The interaction scores of the combinations chosen for testing are shown as red dots. The 35 high interest agents contain drugs either currently used to treat TB or have been used in the past to treat TB (27). **(B)** Comparison of INDIGO-MTB interaction scores with experimental *in vitro* interaction scores. Each dot indicates a specific drug combination. Dark red dots mark two-drug regimens (R = 0.62, p = 9.3*10^−3^), and blue dots mark three-drug regimens (R = 0.64, p = 8.81*10^−2^). The specific combinations mentioned in the text are highlighted in the plot. For both experimental and INDIGO-MTB scores, values less than 0.9 indicate synergy, values between 0.9 and 1.1 denote additivity, and values greater than 1.1 indicate antagonism. **(C)** Dot plot of experimentally measured drug interaction scores versus the INDIGO-MTB predicted drug interaction type. The dots labeled in red font denote outlier combinations that were misclassified by INDIGO-MTB. The interaction scores were significantly different between predicted synergistic and antagonistic combinations (p = 0.0009, KS test). The horizontal lines in the box plot represent the median and the first and third quartiles. **(D)** ROC curves plotting sensitivity vs specificity for INDIGO-MTB predictions of synergy and antagonism for both 2-drug and 3-drug combinations in the validation set. Sensitivity measures the true positive rate, which is the fraction of true positive interactions correctly identified; specificity measures the true negative rate. The area under the ROC (AUC) values provides an estimate of the sensitivity and specificity of model predictions over a range of thresholds. The AUC values are 0.89 and 0.91 for synergy and antagonism respectively. (Sensitivity = 90.9% and Specificity = 84.6% for synergy, Sensitivity = 66.6%, Specificity = 91.7% for predicting antagonism).

We first classified experimentally measured combinations as synergistic, additive, or antagonistic. INDIGO-MTB predicted interaction scores were significantly different between these three classes (p-value = 0.0064, Kruskal-Wallis Rank Sum Test, **Figure 2C**). In addition, there was a significant difference between INDIGO-MTB predictions for synergistic and antagonistic combinations (p-value = 0.0009, non-parametric Komolgorov-Smirnov test). Receiver Operating Curve (ROC) analysis of INDIGO-MTB predictive performance yielded an area under the curve (AUC) of 0.89 (p = 1.2 × 10^−3^) and 0.91 (p = 6.7 × 10^−4^) for detecting synergy and antagonism in the validation set, respectively (**Figure 2D**). These results are robust to the choice of thresholds used for classifying interactions as synergistic or antagonistic (**Figure S4**). We next performed a quantitative comparison between INDIGO-MTB interaction scores and the corresponding *in vitro* experimentally measured FIC indices using the scale invariant metric, spearman’s rank correlation (R) (**Figure 2B)**. We observed a high degree of correlation between model prediction and experimental measurements for all combinations (R = 0.63, p = 9.5 × 10^−4^), and also after separating pairwise (R=0.62 ± 0.03, p = 9 × 10^−3^) and three-way interactions (R=0.64 ± 0.1, p = 8 × 10^−2^). The correlation with INDIGO-MTB predictions is identical for both the novel set (rank correlation R = 0.63) and for the total validation set, (R = 0.64 for all 36 combinations). Thus, not only can INDIGO *qualitatively* differentiate synergy and antagonism, but it can also *quantitatively* separate regimens based on their extent of synergy.

Of note, we validated the INDIGO-MTB prediction that the combination of moxifloxacin (MXF) and spectinomycin (SPC) are pairwise-antagonistic (DiaMOND FIC = 1.50) but could be made more synergistic with the addition of clofazimine (CFZ) (DiaMOND FIC = 0.14). The synergy identified between capreomycin (CAP) and CFZ (DiaMOND FIC = 0.70) and strong antagonism between STM and moxifloxacin (MXF) (FIC = 3.68) were also experimentally confirmed. These results, along with tenfold cross validation analysis of the training data (**Figure S4**), show that INDIGO-MTB can successfully infer novel interactions among drugs with known transcriptome profiles.

While most predictions were confirmed experimentally, there were systematic inconsistencies between the model and experiment for some individual drugs. For example, half of the inconsistencies arose in combinations featuring spectinomycin (SPC). Although SPC has been found to synergize with several anti-TB drugs with multiple modes of action (28, 29), the model tends to overpredict synergy for combinations that include SPC. This may be in part because SPC predictions were based on chemogenomic data from *E. coli* rather than MTB response transcriptomes.

Given the high accuracy of our model for both pairwise and multi-drug combinations, we inferred interactions for 35 promising TB drugs using INDIGO-MTB. The resulting compendium of 6545 three-way, 52,360 four-way, and the top 100 synergistic and antagonistic combinations from 324,632 five-way combinations is provided as a supplement to serve as a resource for guiding future drug combination screens (**Table S2, Table S3**).

### *In vitro* drug synergy is correlated with a surrogate marker of clinical efficacy

We next tested if *in vitro* drug interaction outcomes would be predictive of clinical efficacy. A systematic evaluation of the clinical relevance of *in vitro* drug interactions on treatment efficacy is lacking (30). We therefore compared INDIGO-MTB *in vitro* drug interaction predictions with a meta-analysis of data assembled from 57 phase 2 clinical trials (31). These trials reported regimen efficacy outcomes by sputum culture conversion rates of TB patients at two months. If separate clinical studies reported conflicting efficacy scores for a drug regimen, we used both values for comparison with INDIGO-MTB to incorporate this uncertainty.

We found a highly significant degree of correlation between the INDIGO-MTB interaction scores and the sputum culture conversion rates for the corresponding combinations (R = −0.55 ± 0.04, p ∼ 10^−5^, see **Figure 3A, Table S1D**). The results show that regimens predicted to have greater synergy performed better in the clinical trials. For example, the INH-RIF-STM regimen (green) was predicted to be synergistic *in vitro*, and this combination conferred high patient culture negativity (∼94%) at two months (**Figure 3A**). In contrast, the pairwise combinations of INH-STM (yellow) and INH-RIF (pink) were identified as antagonistic, and both drug pairs resulted in low sputum conversion rates. There was a highly significant difference in sputum conversion between synergistic and antagonistic combinations (p ∼ 10^−4^, **Figure 3B**), the difference in clinical outcome for synergistic-additive (p = 0.038) and additive-antagonism (p = 0.016) interactions were significant as well.

**Figure 3.**
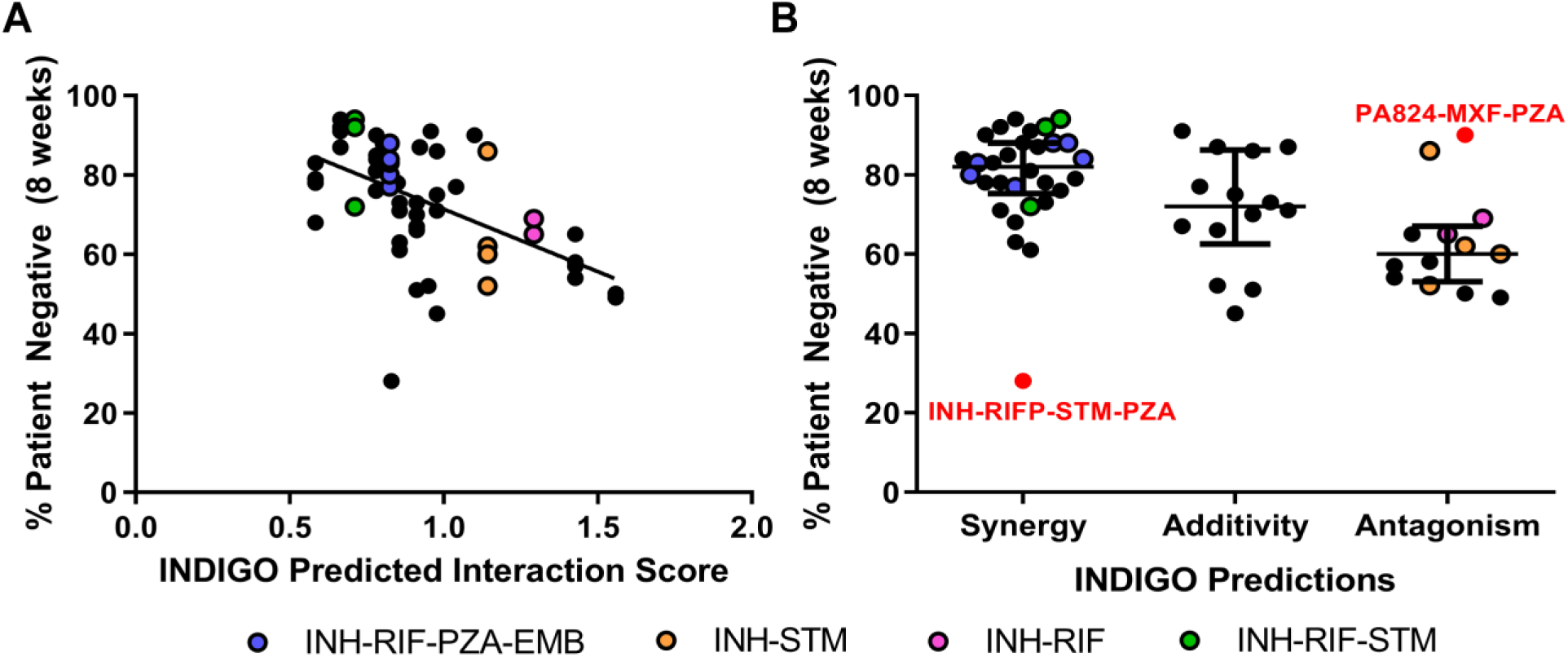
INDIGO-MTB drug interaction scores correlate with sputum culture negativity at 2 months. **(A)** Comparison of model predictions with sputum conversion rates in human patients after 8 weeks of treatment in clinical trials (R = −0.55, p ∼ 10^−5^). Higher patient negative percentages indicate more effective regimens. Each dot indicates a specific drug combination reported from a specific clinical trial. Dots highlighted in the legend are drug combinations of interest mentioned in the text. **(B)** Dot plot of sputum conversion rates against the INDIGO-MTB predicted drug interaction type. The dots labeled in red font denote outlier combinations that were misclassified by INDIGO-MTB. The horizontal lines represent the 1st quartile, 3rd quartile, and median (the widest horizontal line). The colored dots correspond to combinations highlighted in the legend.

Among the combinations assessed clinically, only four two-way and two three-way drug combinations had experimental *in vitro* drug interaction data. We next compared the correlation of *in vitro* experimentally measured drug interaction with the corresponding sputum conversion rates. We found that *in vitro* experimental drug interaction scores also correlated significantly (R = −0.52 ± 0.1, p = 0.01) with clinical sputum conversion by sampling analysis (**Figure S5**). This correlation is comparable to the value observed with INDIGO-MTB across all 57 clinical trials.

Despite the strong overall concordance between *in vitro* synergy and *in vivo* sputum culture conversion rates, we found some outlier combinations that were inferred to be synergistic but had poor clinical outcomes. All the outlier regimens contained pyrazinamide (PZA), whose interaction scores were estimated based on transcriptomes that were generated under acidic conditions, which were unlike the conditions of the other drug profiles. Furthermore, the RIF-MXF combination was identified to be antagonistic by both our model and experiments but has good *in vivo* efficacy. It is hypothesized to be effective because of its ability to suppress resistance despite being antagonistic (32). Hence, synergy alone does not always imply clinical efficacy. Numerous other factors can impact treatment outcome. Combinations can perform well despite being antagonistic. Overall, our results suggest that drug synergy is significantly correlated with treatment efficacy at 8 weeks, and identifying synergistic drug interactions is a promising strategy to prioritize combination regimens.

### Inferring molecular mediators of drug synergy

To interrogate what molecular processes underlie INDIGO-MTB’s predictive ability, we identified genes in the INDIGO-MTB model that most strongly influenced drug interaction scores. Genes were *in silico* “deleted” from the INDIGO-MTB model (i.e., excluded from the model prediction) and assigned an importance score by INDIGO-MTB proportional to their relative contribution in calculating drug interaction scores. The top 500 genes sorted based on their importance score accounted for 97% of INDIGO-MTB’s predictive ability. We performed pathway enrichment analysis using literature-curated pathways from the KEGG database (33, 34) to determine over-represented pathways among the top 500 informative genes (**Table S1E**). Metabolic pathways were highly enriched overall, and the most overrepresented pathway was oxidative phosphorylation, which is targeted by BDQ. The model thus suggests that targeting this pathway might have an impact on drug interaction outcomes.

We hypothesized that we could gain further insights into the genetic regulation of drug interaction outcomes. To do this, we analyzed the INDIGO-MTB model in the context of the MTB transcriptional regulatory network (TRN). The TRN was reconstructed by transcriptome profiling of a comprehensive library of transcription factor induction strains (TFI)(35, 36). The regulon (i.e., set of functional targets) for each transcription factor (TF) was defined as those genes that significantly changed expression upon chemical induction of the TF expression.

To assess the systems-level impact of each TF on drug interactions, we performed *in silico* deletions of entire regulon-defined gene sets and assessed the effect on the INDIGO-MTB interaction scores. We identified regulon deletions that disrupt a specific drug interaction and those that influence multiple drug interaction outcomes. For this analysis, we considered all 36 pairwise combinations comprising the drugs: INH, RIF, STM, MXF, CFZ, BDQ, capreomycin (CAP), ethionamide, and pretomanid (PA824). The drugs tested are all current first- and second-line TB agents that can be prescribed together as part of therapy and have differing mechanisms of action. From this analysis, INDIGO-MTB identified the transcription factor Rv1353c as having the highest impact on drug interactions among all the TFs (**Figure S6A**). INDIGO-MTB estimated that Rv1353c would shift the interaction scores for almost every pairwise interaction toward synergy upon induction (Δscore = −0.6±0.1). The exception was the combination CFZ-STM, for which INDIGO-MTB predicted minimal interaction shift associated with TF induction (Δscore = −0.2) (**Figure S6B**).

We tested these model predictions by comparing the interactions of three representative drug combinations with the following three genetic perturbations: (1) TF induction, measured in the TFI strain with the presence of chemical induction; (2) TF disruption, measured in a knockout strain (see Methods); and (3) baseline TF levels, measured in the genetic wildtype strain, H37Rv and the TFI strain in the absence of chemical induction. We selected two drug combinations for which strong interaction shifts were inferred upon TF induction (BDQ-STM, Δscore = −0.7; CAP-STM; Δscore = −0.7), as well as the CFZ-STM combination for which the model estimated minimal interaction shift. The baseline interactions between the drug combinations differ substantially (BDQ-STM is additive, whereas CAP-STM and CFZ-STM are both antagonistic, **Figure S6C**). **Figure 4** shows the difference in experimentally measured interaction scores of each drug combination for the genetic perturbation conditions, relative to the wildtype (Methods). The results show that when Rv1353c is induced, interactions for both BDQ-STM and CAP-STM shift toward synergy (ΔFIC = −0.2 ± 0.1, p = 0.03 for BDQ-STM; ΔFIC = −0.5 ± 0.2, p = 0.01 for CAP-STM), and when Rv1353c is disrupted, interactions for both BDQ-STM and CAP-STM shift toward antagonism (ΔFIC = 0.3 ± 0.2, p = 0.001 for BDQ-STM; ΔFIC= 0.2 ± 0.2, p = 0.04 for CAP-STM). In contrast, there appears to be no significant shifts in interaction for CFZ-STM with either induction or disruption of Rv1353c (ΔFIC = −0.0004 ± 0.3, p = 0.5 for disruption; ΔFIC= −0.03 ± 0.1, p = 0.03 for induction). Collectively, these results confirm the INDIGO-MTB predictions.

**Figure 4.**
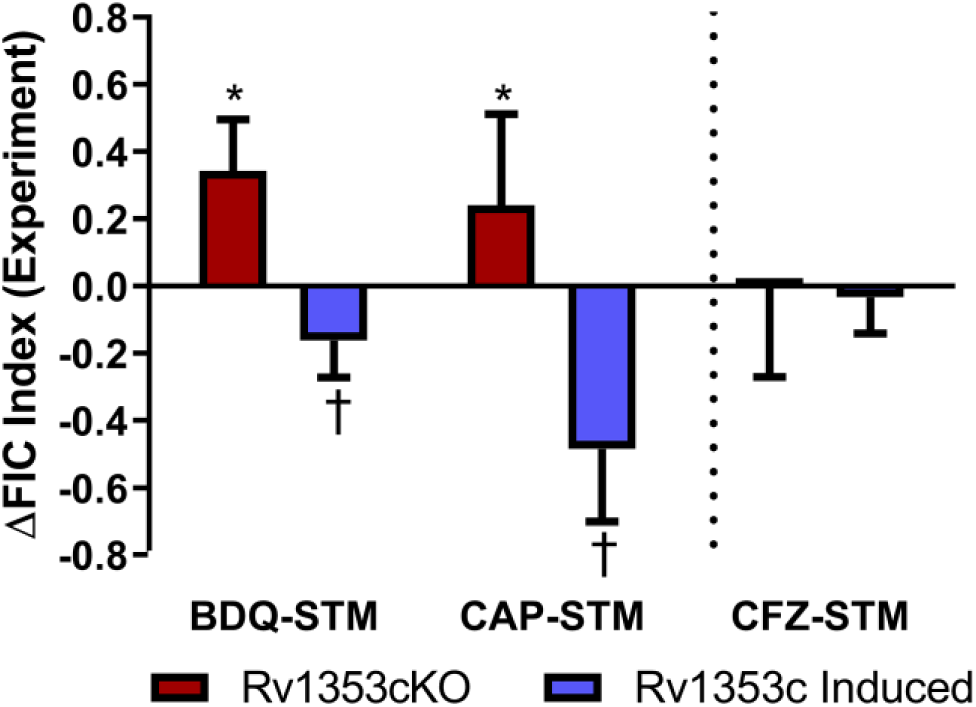
Rv1353c influences interactions between drug combinations. The *in vitro* experimentally measured drug interaction scores are quantified for the three selected drug interactions, plotted as the difference in FIC score of the gene perturbation relative to the wildtype (H37Rv). The red bars denote values for the knockout strain, and the blue bars show values for the strain with Rv1353c induced. Negative values indicate shifts toward synergy, and positive values indicate shifts toward antagonism. The (*) and (†) indicate that differences are significantly greater or less than zero, respectively (p < 0.05, one-tailed one-sample t-test). The error bars represent the standard deviation between replicates.

## DISCUSSION

Here, we constructed an INDIGO-MTB model to predict *in vitro* synergy and antagonism of anti-tuberculosis drug combinations using transcriptomics data. Our model complements existing experimental strategies by increasing throughput and by identifying potential drug interaction mechanisms. Our analysis using INDIGO-MTB revealed novel synergy between clinically promising drug combinations, uncovered the role of the TF Rv1353c in influencing drug interaction outcomes, and found a significant association between *in vitro* drug interaction outcomes and clinical efficacy. These results suggest that using INDIGO-MTB to identify synergistic regimens is a promising strategy for prioritizing combination therapies. While significant challenges exist, constructing a high-quality model of drug interactions *in vitro* is the first step towards inferring *in vivo* efficacy. No theoretical method currently exists that can comprehensively screen thousands of combinations even *in vitro*. The significant correlation between INDIGO interaction scores with both *in vitro* data and clinical efficacy data supports the utility of our approach.

INDIGO-MTB outperforms existing strategies in terms of throughput. The largest studies in MTB have so far analyzed up to two hundred unique drug combinations (37). Here, we have estimated outcomes for 13,366 pairwise and 721,764 three-way combinations of 164 drugs with significant accuracy based on our prospective validation. While many of the drugs might have poor anti-TB activity on their own, they may greatly enhance synergy when added to existing regimens. For example, we found chlorpromazine, originally used for treating psychiatric disorders, synergizes with BDQ, resulting in four-fold reductions in inhibitory concentrations (**Figure S7A**). Thus, INDIGO can facilitate repurposing of drugs to treat TB.

INDIGO complements other preclinical methods such as mouse models in prioritizing regimens for clinical evaluation. A systematic comparison across multiple mouse studies is challenging due to the lack of quantitative raw data and variation in metrics reported in the literature. Nevertheless, combinations identified by INDIGO to be highly synergistic (top 0.01%, **Table S2, Table S3**) were also found to be highly efficacious in recent mouse studies. Combinations involving BDQ and CFZ alone or in a three-drug combination with PZA, Ethambutol (EMB), RIF, or INH were all found to be synergistic by INDIGO and showed high bactericidal activity in mouse models (5, 38-40). Four-way drug combinations involving BDQ, CFZ and PZA with EMB or SQ109 were also synergistic in mouse studies (5, 38-40). In addition to these combinations studied in mouse models, INDIGO-MTB also uncovered highly synergistic novel 4-drug and 5-drug combinations that are promising candidates for pre-clinical evaluation, such as the combination with BDQ, CFZ, RIF, CLA and the anti-malarial antifolate compound P218, and a 4-drug combination involving BDQ, RIF, PA824, and the anti-psychotic drug thioridazine (**Table S3**).

Since numerous factors could impact *in vivo* efficacy that are not considered during *in vitro* studies, it is not *a priori* clear if there should be a significant correlation between *in vitro* synergy and *in vivo* efficacy. Thus, we performed a systematic comparison of *in vitro* drug interaction scores with clinical efficacy of drug combination regimens. Notably, here we observed a statistically significant correlation between *in vitro* drug interaction scores and the percentage of TB patients showing negative sputum culture after 2 months treatment in clinical trials, with synergistic drug combinations showing greater clinical efficacy. Negative sputum culture at eight weeks is a useful early measure of TB treatment efficacy that correlates well with relapse rates (41, 42). The correlation that we observed between *in vitro* INDIGO-MTB predictions and sputum conversion rates is notable, given the huge variability between clinical studies.

While existing high-throughput approaches are strictly non-mechanistic, INDIGO can reveal the relative contribution of underlying cellular pathways on drug interaction outcomes. Our analysis suggests that drug transporters and central metabolic pathways may play a role in influencing drug interaction outcomes. This is consistent with recent studies on the role of bacterial metabolic state in impacting drug interaction outcomes (43, 44). Contextualizing INDIGO-MTB with the MTB transcriptional regulatory network revealed genetic regulators of drug interaction response. This analysis uncovered the role of the transcription factor, Rv1353c as a broad regulator of drug interaction outcomes. Rv1353c is an uncharacterized nonessential helix-turn-helix type transcriptional regulator (45-49) that has previously been found to be deleted in several clinical isolates (50). When induced under log-phase growth, Rv1353c activates 44 genes enriched for fatty acid biosynthesis and represses 50 genes, including two of the top five most informative INDIGO-MTB predictor genes (Rv1857 and Rv1856c)(35). Interestingly, INDIGO-MTB simulations suggest minimal shifts in drug interaction scores upon perturbing either Rv1857 or Rv1856 individually, suggesting that the underlying molecular mechanisms mediating drug interactions may be partially epistatic in nature. Collectively, this suggests that knowledge of the underlying mechanism of drug interaction can be used to engineer synergy between combination regimens. Our approach provides a rational strategy to identify genetic targets that enhance synergy between existing regimens and introduces a potentially new way to engineer effective regimens by modifying the interactions between the constituent drugs.

While the INDIGO approach has demonstrated significant utility in predicting synergy and antagonism of drug combinations, it nevertheless has several key limitations. First, INDIGO-MTB requires as input transcriptome data profiling of MTB response to each drug for which drug interaction predictions are necessary. Transcriptomes are significantly faster and cheaper to generate than the chemogenomic profiles used to power the original INDIGO models. This has enabled us to use species-specific data to build INDIGO-MTB. Among the 35 TB drugs of interest, the input data for only 10 drugs (28%) are derived from *E. coli* chemogenomics data. The correlation observed in the current study, wherein the model was constructed using MTB response transcriptomes elicited by drug exposure, is higher than the correlation observed in our prior study, which used chemogenomic data to infer interactions (R = 0.62 for pairwise and 0.64 for three-way interactions for the current study, versus R = 0.52 for pairwise and 0.56 for three-way interactions in the *E. coli* chemogenomic study (14, 51)). Notably, while predictions using *E. coli* data were statistically significant, many of the incorrect predictions from our model, such as drug combinations involving spectinomycin, might be attributed to challenges of extrapolating predictions from *E. coli* using gene orthology information alone. Our results suggest that gene expression changes encapsulate molecular response information that is as informative of drug interaction phenotypes as gene deletion studies. The updated INDIGO approach can hence be applied to other pathogens that lack chemogenomic data. With reduced sequencing costs, transcriptomics data is unlikely to be a substantial limitation in the future. Further, while the number of possible combinations increases exponentially with the number of drugs, the number of transcriptomes required only increases linearly. Hence, INDIGO-MTB and other methods that use responses elicited by individual drugs will be more cost and time effective.

A second limitation stems from the fact that INDIGO-MTB predictions are currently based on data gathered from log-phase *in vitro* broth culture conditions, which are markedly different from the *in vivo* microenvironments. Outliers from our experimental validation involving PZA (which is relatively more active under low pH conditions) substantiate the notion that the underlying environmental context can influence the model accuracy. The INDIGO algorithm is currently blind to MTB molecular responses to drugs in the host context. Training our model using MTB transcriptome profiling data generated using an appropriate environmental condition (e.g., MTB in a macrophage or mouse infection model) might address this limitation in the future. A recent study has expanded the INDIGO model to enable *in silico* prediction of the impact of different microenvironments in *E. coli* (51). Hence building an accurate INDIGO model for MTB can provide a foundation for addressing this *in vivo* complexity.

Finally, while synergy is associated with a better treatment outcome on average, other factors such as resistance evolution, toxicity, and drug pharmacokinetics will also influence treatment success. In addition, there is considerable heterogeneity in clinical trial efficacy based on patient population, dose and location. The curation of numerous clinical studies and ability to predict interactions in high throughput provided us with sufficient statistical power to test the association between synergy and *in vivo* efficacy despite this heterogeneity. In the future, incorporating additional factors associated with drug behavior in the host may further improve the correlation between model predictions and clinical outcomes.

## METHODS

### Culture conditions

MTB strains were cultured in Middlebrook 7H9 with the oleic acid, bovine albumin, dextrose and catalase (OADC) supplement (Difco), and 0.05% Tween80 at 37 °C under aerobic conditions with constant agitation to mid-log phase, as described previously (35, 52). Strains containing the anhydrotetracycline (ATc)-inducible expression vector were grown with the addition of 50 μg/mL hygromycin B to maintain the plasmid. To induce expression of the transcription factor Rv1353c, 20ng/uL of ATc was added to the culture media. Growth was monitored by the optical density at 600 nm (OD600).

The Rv1353c overexpression strain was generated previously (35, 36). Briefly, the Rv1353c gene was cloned into a tagged, inducible vector that placed the gene under control of a tetracycline-inducible promoter (53) and added a C-terminal FLAG epitope tag. This construct was transformed into MTB H37Rv using standard methods. The strain is available from the BEI strain repository at ATCC ((54), NR-46512).

### Phage Knockout Strain Generation

The H37Rv Δ*Rv1353c* strain was constructed by a specialized transduction method(55) using a gene-specific specialized transducing phage phasmid DNA provided by the Jacobs lab and the previously described protocol (55). Briefly, high-titer phage stocks were generated by transfecting the phasmid DNA into *Mycobacterium smegmatis* mc^2^155 at 30°C, and growing the resulting phage plaques on an agar pad with a lawn of mc^2^155. Transduction-competent H37Rv was incubated with high-titer phage stock for 24 hours at 37°C, and the transduced bacteria were plated on 7H10 supplemented with 50 μg/mL hygromycin B to select for deletion-substitution mutants.

### Drug susceptibility and checkerboard drug-drug interaction experiments

Strains were grown to log phase (OD600 ≈ 0.3), diluted to a final OD600 ≈ 0.005 (equivalent to 10^6^ (CFU)/mL), and dispensed into 96-well flat-bottom plates (Corning, Acton, MA) at a final volume of 200µL, containing 1% DMSO and varying concentrations of drugs in the different wells. On each plate, control wells for each of the strains studied were included, containing: 1) no drug and 1% DMSO vehicle; and 2) 1% culture and no drug with 1% DMSO vehicle, to measure viability in the absence of drug exposure.

For drug susceptibility assays to measure the MIC, serial 2-fold dilutions of an individual drug were arrayed in the different columns. For checkerboard drug interaction assays, 2-fold dilutions of the first drug were arrayed in the columns and 2-fold dilutions of a second drug were arrayed in the rows.

Plates were incubated at 37°C for 7 days. Cellular viability was assayed on day 7 by the BacTiter Glo (Promega, Madison, WI) and Alamar Blue cell proliferation assays (Bio-Rad, Hercules, CA) according to manufacturer recommendations. Briefly, we added 20µL of culture from each well to 20µL of BacTiter-Glo Microbial Cell Viability Assay Reagent, incubated at room temperature protected from direct light for 20 minutes, and read luminescence intensity using a FluoStar Omega plate reader (BMG Lab Tech, Cary, NC). For Alamar Blue, we added 20µL of Alamar Blue reagent to 180µL of culture, incubated for 12 hours protected from direct light, and read fluorescence intensity at emission wavelength 590nm after excitation at 544nm. **Figure S7B** shows the strong concordance between the two methods - BacTiter-Glo and Alamar Blue.

For drug susceptibility assays, the MIC was determined as the lowest drug concentration that resulted in MTB viability comparable to the 1% culture control. For checkerboard assays, the drug interaction was quantified by the Fractional Inhibitory Concentration (FIC) index, equal to: 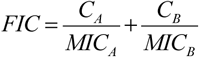, where C_A_ is the concentration of drug A when combined with drug B yielding an iso-effective inhibition comparable to the MIC, and C_B_ is the concentration of drug B when combined with drug A yielding an iso-effective inhibition. The value for FIC can be extended to any arbitrary number of drug combinations as follows

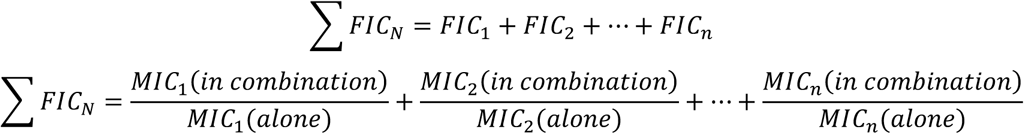

Each MIC and checkerboard experiment was performed 2 times, with 2 biological replicates per experiment. The mean FIC index across all iso-effective concentrations was calculated for each biological replicate to determine reproducibility, and data across biological replicates were summarized by averaging (**Figure S1**).

### DiaMOND drug-drug interaction experiments

DiaMOND drug interaction experiments were performed in biological triplicate as previous described (6). Rather than sampling the entire set of dose combinations used in a traditional checkerboard assay, DiaMOND samples a subset of dose responses and approximates the shape of the contour of the chosen phenotype (e.g. where 50% growth inhibition is observed, IC50). For example, a two-drug combination requires three dose responses (each individual drug dose response and an equipotent drug combination dose response) rather than the entire set of possible dose combinations.

Individual drug dose response ranges were chosen for each drug such that the IC50 dose was close to the center and doses were linearly spaced to provide high resolution IC50 determination. Drug combination dose response ranges contained equipotent mixtures of two or three drugs (e.g. a two-drug combination would contain ½ of the IC50 dose for each drug and a three-drug combination would contain ⅓ of the IC50 of each drug).

Briefly, MTB strain H37Rv cultures were grown to mid-log phase (OD600 ≈ 0.6), diluted to OD600 ≈ 0.05 and added to drug containing plates. Drugs were dispensed into 384-well plates using a digital drug dispenser (D300e Digital Dispenser, HP) and 50 μL diluted MTB cultures were overlaid. Drug treatment plates were incubated in humidified containers for 5 days at 37 °C without agitation. Growth was measured by OD600 using a plate reader (Synergy Neo2, Biotek). Two technical replicates were performed, and the average of each technical replicate was used to calculate FIC scores.

The FIC for a drug combination was calculated as the ratio between the observed and expected IC50 dose of the drug combination as previously described (6). FICs from each of three biological replicates were calculated to determine reproducibility, and data across biological replicates were summarized by averaging. Briefly, the growth measurements were normalized (background subtracted, normalized to untreated) and the observed IC50 doses were calculated for each individual and combination drug dose response. The expected IC50 dose for the drug combination was then calculated using the IC50 of the individual drugs, based on the null hypothesis that the interaction is additive. For two-drug combinations the expected IC50 dose is defined as the intersection of the line (additivity line) drawn between the IC50 doses for each individual drug. For three-drug combinations, the expected IC50 dose is defined as the intersection of the drug combination dose response and the plane (additivity plane) created by connecting the IC50 doses for each individual drug (**Figure S1**).

### RNA-seq transcriptome profile data generation

To profile the MTB transcriptome response to exposure of individual drugs, cultures were diluted to OD600 ∼ 0.2 (equivalent to 10^8^ colony-forming units (CFU)/mL) and exposed to a minimum inhibitory concentration (MIC)-equivalent dose of drug for approximately 16 hours.

RNA was isolated from these cultures as described previously (35, 52). Briefly, cell pellets in Trizol were transferred to a tube containing Lysing Matrix B (QBiogene) and vigorously shaken at maximum speed for 30 s in a FastPrep 120 homogenizer (QBiogene) three times, with cooling on ice between shakes. This mixture was centrifuged at maximum speed for 1 min and the supernatant was transferred to a tube containing 300 μL chloroform and Heavy Phase Lock Gel (Eppendorf), inverted for 2 minutes and centrifuged at maximum speed for 5 minutes. RNA in the aqueous phase was then precipitated with 300 μL isopropanol and 300 μL high salt solution (0.8 M Na citrate, 1.2 M NaCl). RNA was purified using a RNeasy kit following the manufacturer’s recommendations (Qiagen) with one on-column DNase treatment (Qiagen). Total RNA yield was quantified using a Nanodrop (Thermo Scientific).

To enrich the mRNA, ribosomal RNA was depleted from samples using the RiboZero rRNA removal (bacteria) magnetic kit (Illumina Inc, San Diego, CA). The products of this reaction were prepared for Illumina sequencing using the NEBNext Ultra RNA Library Prep Kit for Illumina (New England Biolabs, Ipswich, MA) according to manufacturer’s instructions, and using the AMPure XP reagent (Agencourt Bioscience Corporation, Beverly, MA) for size selection and cleanup of adaptor-ligated DNA. We used the NEBNext Multiplex Oligos for Illumina (Dual Index Primers Set 1) to barcode the DNA libraries associated with each replicate and enable multiplexing of 96 libraries per sequencing run. The prepared libraries were quantified using the Kapa qPCR quantification kit, and were sequenced at the University of Washington Northwest Genomics Center with the Illumina NextSeq 500 High Output v2 Kit (Illumina Inc, San Diego, CA). The sequencing generated an average of 75 million base-pair paired-end raw read counts per library.

Read alignment was carried out using a custom processing pipeline that harnesses the Bowtie 2 utilities(56, 57), which is available at https://github.com/sturkarslan/DuffyNGS, and https://github.com/sturkarslan/DuffyTools. The RNA-seq data profiling response to drug exposure generated for this study are publicly available at the Gene Expression Omnibus (GEO) at **GSE119585**.

### Gene expression data analysis

The RNA-seq transcriptome profiling data that we generated were supplemented with microarray and RNA-seq transcriptome profiling datasets from literature that were downloaded from GEO, along with associated gene accession identifiers. The log_2_-transformed fold change values of average gene expression in each treatment group were determined for all studies, relative to the experiment’s negative control. All genes that significantly change by more than 2-fold (up or down) after each drug treatment were used as input features for INDIGO-MTB. The results are robust to the thresholds chosen for finding differentially expressed genes (**Table S1G**).

ComBat (18) normalization was used to minimize batch effects in the data, which uses empirical Bayes approach to estimate each batch’s corrected mean and variance. The effectiveness of normalization was checked using principal component analysis. This version of the transcriptomic/chemogenomic matrix represented the drug-gene network that was required to build the INDIGO-MTB model.

The drug-gene interaction profiles for each drug are then used by INDIGO to create a “joint” interaction profile for a drug combination (**Figure S1**). INDIGO assumes that cellular response to drug combinations is a linear function of the cellular response to individual drugs. This assumption is based on prior experimental studies that found that a linear model best explained transcriptional response of cells treated with drug combinations (58, 59). Further, in our prior study in *E. coli*, we found that other models of profile integration, such as correlation or profile overlap performed poorly in predicting drug interactions compared to the linear integration model (14).

### Quantifying drug-drug interaction scores for model training

To train INDIGO-MTB, checkerboard FIC indices of drug combinations were collected after conducting literature search (n=140). We also included FIC50 indices that were calculated using the DiaMOND approach (n=62)(6). Since the DiaMOND study had a distinct distribution from other checkerboard studies from literature (Mean = 1.05 and 0.99, Standard deviation = 0.32 and 0.81 for DiaMOND and checkerboard respectively), we statistically transformed the DiaMOND scores so that the overall distribution of the DiaMOND-measured scores had the same mean and standard deviation as the remaining checkerboard datasets. The normalized scores were used for training INDIGO-MTB (**Table S1F**). Similarly, the DiaMOND data generated in this study for validation was normalized using the same approach prior to comparison with INDIGO-MTB predictions. The average interaction score in the final training set was 1.01, suggesting that the training data set is not significantly biased towards synergy or antagonism.

### Statistical Analyses

Our experimental test set is sufficiently powered statistically to significantly assess the accuracy of INDIGO’s correlation with the experimental data. For example, the probability of getting a correlation of 0.62 achieved by INDIGO by random chance is less than 1 in 10^3^. We statistically estimated that we only need 14 samples to detect a correlation of 0.6 (R>=0.6) with a p-value of 0.01. Our test set sample size is significantly larger than this number.

Spearman rank correlations were computed using the statistical software R. Differences between the means of each group in box plots were compared using two-sample one-tailed K-S tests in R. To further assess the robustness of our results to variation in clinical trials, we performed sampling analysis by choosing one representative clinical trial randomly for each regimen. We observed a significant correlation between predicted interaction scores and the sputum culture conversion rates (mean rank correlation R = −0.38 average of 100 random sampling trials) (**Figure S7B**).

The significance of the AUC values from the ROC analysis was calculated by randomly permuting the class labels (synergy or antagonism) of the test data 1000 times. The difference in accuracy of the actual model with the random permuted models was compared using a t-test.

We used the RandomForest algorithm that is part of the Machine learning toolbox in MATLAB. The regression random forest algorithm was used with default parameters for the number of predictors sampled (default value – N/3, where N is the number of variables). Hyperparameter tuning of parameters in the training set instead of using default parameters also resulted in a similar accuracy in the test set (**Table S1H**). Random forests are perfectly suited for our analysis as they can achieve high accuracy even with small sample sizes and can be easily interpreted. The training set used here is relatively small for deep neural networks which require thousands of samples. On the other hand, SVM and decision trees can be built with small sample sizes but do not achieve high accuracy as Random Forests. The accuracy using these approaches with default parameters is lower than Random Forests with default parameters (**Table S1H**).

The INDIGO-MTB model and associated data sets are available from the Synapse bioinformatics repository (Synapse ID: syn18824984) (https://www.synapse.org/INDIGO_MTB) (DOI: 10.7303/syn18824984).

## ACKNOWLEDGMENTS

The data reported in the paper are available in the Supplementary Materials. The RNA-seq data generated for this study are available on the Gene Expression Omnibus [GSE119585].

We gratefully acknowledge the laboratory of William R. Jacobs Jr. for providing the phage that enabled us to make the Rv1353c knockout strain. We gratefully acknowledge Andréanne Lupien, Anthony Vocat, and Stewart T. Cole for providing us a sample of PBTZ169 for our study. We also gratefully acknowledge the laboratory of Timothy Sterling for providing us with ofloxacin resistant clinical strains. We also thank Jessica Winkler, Laura Green, and Reiling Liao, for their technical assistance, and Awanti Sambarey for her feedback on the manuscript.

This work was supported by the National Institutes of Health [grant numbers U 19 AI106761 (DS, SC); U19 AI111276 (DS); U19 AI135976 (DS); 5T32AI007509 (SM); P50 GM107618-01A1 (BA); UL1TR002240 (SC); NIH Director’s New Innovator Award 1DP2LM011952-01(BA); the University of Michigan Precision Health and MCUBED (SC).

## AUTHOR CONTRIBUTIONS

S.M. conceived of the study, led the design, generated experimental checkerboard and RNA-seq data, analyzed the experimental data, and drafted the manuscript. S.J. co-developed the INDIGO-MTB model, generated the model predictions. J.L-F. generated DIAMOND experimental drug interaction data and drafted the manuscript. J.L. generated experimental checkerboard drug interaction data and RNA-seq data. B.A. organized the DIAMOND experimental drug interaction data generation and drafted the manuscript. D.S. and SC conceived of the study, led the design, organized the data analysis, and drafted the manuscript, and SC also co-developed the INDIGO-MTB model.

## SUPPLEMENTAL MATERIALS

**Figure S1**: **Schematic of INDIGO-MTB modeling workflow (A-C) and the experimental assays to measure drug interaction (D-F). (A)**The input datasets for training the INDIGO algorithm are: [1] the transcriptomic profiles of drugs, and [2] the corresponding FIC scores of known drug-drug interactions. From each transcriptome profile of MTB response to an individual drug, we defined a corresponding drug-gene interaction matrix by assigning a value of 1 to genes that changed in expression by more than 2-fold (up or down) after exposure to drug, and setting all other genes to a value of 0. Only genes that are up-regulated are shown in the remaining panels for simplicity. **(B)** For each drug combination, INDIGO calculates a “joint” drug-gene interaction matrix using a linear combination of the drug-gene interaction matrices of each constituent individual drug. The joint profile captures both the similarity and uniqueness in the transcriptome response profiles of the individual drugs in each combination. The INDIGO algorithm then uses a machine learning approach called Random Forest to create a mathematical model that associates the FIC score of each drug combination to its corresponding joint drug-gene interaction matrix. Random Forest builds a series of decision trees to identify specific patterns in the drug-gene interaction matrices that significantly associate with the value of the corresponding drug-drug interaction FIC scores. **(C)** Once built, the INDIGO-MTB model requires only the transcriptomic response profile elicited by a new compound of interest as input to predict FIC scores of combinations featuring the compound of interest. **(D)** Representative checkerboard assay experiments of a synergistic and antagonistic drug pair. Cultures were exposed to serial dilutions of drugs (designated in the rows and columns) for 7 days, and bacterial viability was quantified by measuring ATP levels with the BacTiter Glo reagent. The thick black boxes denote the individual drug MIC wells, and the boxes with numbers denote concentrations that yielded iso-equivalent inhibition (each of the numbers represent the FIC score calculated based on the drug concentrations associated with corresponding well). **(E-F)** Representative DiaMOND assay experiments of a synergistic and antagonistic drug pair **(E)** or triplet **(F)**. Cultures were exposed to drugs in 384-well plates and growth was measured by OD600. The FIC for a drug combination was calculated as the ratio between the observed and expected IC50 dose of the drug combination. For two-drug combinations the expected IC50 dose is defined as the intersection of the line (additivity line) drawn between the IC50 doses for each individual drug. For three-drug combinations, the expected IC50 dose is defined as the intersection of the drug combination dose response and the plane (additivity plane) created by connecting the IC50 doses for each individual drug.

**Figure S2. Distributions of INDIGO-MTB interaction scores for combinations featuring different drugs. (A)** Box plots of interaction scores of combinations featuring only bacteriostatic drugs (blue) is shifted toward synergy (interaction score = 1.03±0.2), relative to combinations featuring bactericidal drugs (red) (interaction score = 1.25±0.3). Combinations featuring bactericidal drugs appear to have the most antagonistic INDIGO-MTB interaction scores. **(B)** Distribution of INDIGO-MTB interaction scores for combinations involving Verapamil (VER) or Chlorpromazine (CPZ). The distribution of interaction scores for these drugs is significantly lower than the interaction score distribution for combinations featuring other drugs (p-value < 1 × 10^−16^, non-parametric Kolmogorov-Smirnov test), suggesting that combinations featuring these drugs are enriched for synergy. The box plots display the first quartile (1Q), median, and the third quartile (3Q) of the distribution of INDIGO-MTB scores for pairs of drugs, at least one of which are VER or CPZ compared against all possible combinations excluding these two antibiotics.

**Figure S3: INDIGO accurately forecasts interactions among the 36 test-set drug combinations against MTB.** This figure shows the comparison between INDIGO-MTB interaction scores and experiments for all 36 combinations in the validation set. This figure complements Figure 2 which compares INDIGO predictions with the 24 novel combinations (subset of 36 test combinations). **(A)** Drug combinations chosen for experimental testing span the entire range of drug interaction predictions by INDIGO. The histogram and box plot above it show the distribution of pairwise drug interaction scores for the 35 high interest TB agents (the boundaries of the box plot denote the 25^th^ and 75^th^ percentiles of the distribution, and the dashed lines extend between 1^st^ and 99^th^ percentiles). The red dots denote the combinations selected for experimental validation. **(B)** Comparison of INDIGO-MTB interaction scores with *in vitro* interaction scores. For both experimental and model-predicted scores, values less than 0.9 indicate synergy, values between 0.9 and 1.1 denote additivity, and values greater than 1.1 indicate antagonism. Each dot indicates a specific drug combination. Dark red dots mark two-drug regimens (R = 0.63, p ∼ 10^−4^), and blue dots mark three-drug regimens (R = 0.68, p ∼ 10^−2^). The specific combinations mentioned in the text are highlighted in the plot. **(C)** Dot plot of experimentally measured drug interaction scores versus the INDIGO-MTB predicted drug interaction type. The dots labeled in red font denote outlier combinations that were misclassified by INDIGO-MTB. The interaction scores were significantly different between predicted synergistic and antagonistic combinations (p = 6 × 10^−5^). The horizontal lines in the box plot represent the median and the first and third quartiles. **(D)** Sensitivity vs specificity curves for INDIGO-MTB predictions of synergy and antagonism for both 2-drug and 3-drug combinations in the validation set. The AUC values are 0.89 (p = 4 × 10^−5^) and 0.91 (p = 4.1 × 10^−5^) for synergy and antagonism respectively. (Sensitivity = 88.4% and Specificity = 86.5% for synergy, Sensitivity = 79.4%, Specificity = 90% for predicting antagonism).

**Figure S4: Receiver operating curves (ROC) for INDIGO-MTB predictions of synergy and antagonism for various thresholds of synergy and antagonism. (A-D)** ROC curves generated from the independent test data. Sensitivity measures the true positive rate, which is the fraction of true positive interactions correctly identified; specificity measures the true negative rate. The area under the ROC curve (AUC) values provides an estimate of the sensitivity and specificity of model predictions over a range of thresholds. At the highest threshold for synergy (i.e. FIC < 0.5), the accuracy of the model is reduced likely due to majority of the interactions in the test set being classified as neutral. **(E-H)** ROC curves INDIGO-MTB tested using ten-fold cross validation analysis. In cross validation analysis, 10% of the training dataset is blinded and predictions are made using an INDIGO-MTB model trained using the remaining 90% of the training data. Performance metrics are then calculated based on prediction on the withheld data. This analysis is repeated 10 times to cover the entire training dataset. The plots show the sensitivity vs specificity curves for INDIGO-MTB predictions of synergy and antagonism for various thresholds of synergy and antagonism. The cross-validation accuracy is surprisingly lower than the test set accuracy as the training set comprises data from 15 different studies that were done in diverse labs and batches with different drugs and methodologies. In our prior study (14), the training and test data were obtained from a single source, and consequently the cross-validation accuracy matched the test set accuracy.

**Figure S5. Impact of variation between clinical trials and experimental drug interaction studies on correlation between drug synergy and clinical efficacy.** Since multiple clinical trials and experimental studies had measured the clinical efficacy outcome (sputum conversion rates) and *in vitro* FIC scores for drug combinations, we assessed the robustness of correlations between clinical efficacy data and both experimentally measured **(A-B)** and model-predicted **(C-D)** interaction scores by performing sampling analysis. **(A).** Distribution of the correlation between *in vitro* experimentally measured drug interaction scores with corresponding sputum conversion rates. Since multiple experimental studies had measured the *in vitro* interaction outcome for these combinations, we performed sampling analysis by randomly choosing one representative study for each combination to determine the average correlation. We found that the *in vitro* experimental drug interaction scores correlated significantly (mean R = −0.52, p ∼ 0.01, average of 100 trials) with clinical sputum conversion by this sampling analysis. Panel **(B)** shows one representative trial with R = −0.52. For comparison, INDIGO-MTB achieved a similar correlation across all 57 clinical trials (R = −0.55, p ∼ 10^−5^). **(B)** Comparison of experimental FIC scores with sputum conversion rates in human patients after 8 weeks of treatment in clinical trials, with each regimen represented by FIC data from a single experimental study (R = −0.52, p = 0.01). Data shown for four two-way and two three-way drug combinations that had both experimental *in vitro* drug interaction data and sputum conversion rates. Each dot indicates a specific drug combination reported from a specific clinical trial. The combinations corresponding to each dot is provided in the legend. **(C)** Histogram visualizing the distribution of the correlations between INDIGO-MTB predictions and clinical efficacy, based on sampling analysis. We performed sampling analysis by randomly choosing one representative clinical trial for each combination to determine the average correlation with INDIGO-MTB predicted interaction scores. We observed a significant correlation between interaction scores and the sputum culture conversion rates (mean R = −0.38 average of 100 random sampling trials, p-value = 0.001). **(D)** shows data from one representative trial with R = −0.37.

**Figure S6: Impact of Transcription Factor (TF) deletion on drug interaction scores. (A)** The average predicted impact of deleting each of the 206 TFs on all 36 pairwise combinations comprising INH, RIF, STM, MXF, CFZ, BDQ, CAP, ETA, and PA824 is shown. Rv1353c had the biggest average impact on all the combinations and was chosen for experimental validation. **(B)** Predicted impact of Rv1353c deletion on drug interaction scores. The plot shows the relative absolute difference in FIC scores on all 36 pairwise combinations comprising INH, RIF, STM, MXF, CFZ, BDQ, CAP, ETA, and PA824. Interestingly, combinations with STM showed both the highest and lowest change in interaction score. The interactions in bold were chosen for experimental testing. All the chosen combinations were also antagonistic or additive, thus changing the activity of Rv1353c could be used to make these combinations more synergistic. **(C)** *in vitro* experimentally measured drug interaction scores for Rv1353c genetic perturbation strains exposed to drug combinations. The error bars represent the standard deviation between replicates.

**Figure S7. (A)** *in vitro* experimentally measured interaction score measured between bedaquiline (BDQ) and chlorpromazine (CPZ). The heatmap shows a representative checkerboard assay experiment, in which interaction MTB H37Rv cultures were exposed to different pairwise concentrations of BDQ and CPZ in 96-well plate format, designated in the columns and rows, respectively. Cultures were exposed to drugs for 7 days, and bacterial viability was quantified by measuring ATP levels with the BacTiter Glo reagent. The thick black boxes denote the individual drug MIC wells, and the boxes with numbers denote concentrations that yielded iso-equivalent inhibition (each of the numbers represent the FIC score calculated based on the drug concentrations associated with corresponding well). **(B)** Correlation analysis of FIC average values calculated from checkerboard assays measured by AlamarBlue or BacTiter Glo. Each point represents the average of the FIC indices of equivalently inhibited concentrations on an individual checkerboard plate. Pearson’s R for this is .78 (p < 0.0001).

**Table S1: (A)** Compendium of transcriptomics/chemogenomic data for INDIGO-MTB. **(B)** Training combinations with associated FIC interaction indices. **(C)** List of all possible pairwise interaction scores (164 compounds and 65 perturbations). **(D)** INDIGO-MTB predictions along with corresponding experimental (*in vitro*) and clinical validation data. Highlighted combinations are novel i.e. not used in training set. **(E)** Pathway enrichment analysis using the top 500 predictive genes. **(F)** DiaMOND Distribution Transformation**. (G)** Impact of various thresholds for finding differentially expressed genes after drug treatment. Changing the log fold change threshold from 4-to 32-fold showed similar correlation with the experimental test set interactions as the default threshold (2-fold). The use of a fold change threshold to binarize the data was performed to reduce the noise in the datasets. Overall, this analysis shows that the INDIGO model is robust to the thresholds used. At the highest threshold (32-fold), a slightly higher correlation was observed, although it was not significantly better than the default settings based on partial correlation analysis (p-value > 0.05). **(H)** Impact of using various machine learning algorithms for predicting drug interactions.

**Table S2: 2-way, 3-way, INDIGO-MTB scores for 35 high-interest TB drugs.**

**Table S3: 4-way, and 5-way INDIGO-MTB scores for 35 high-interest TB drugs.**

